# The role of mating systems in postzygotic reproductive isolation between two recently divergent *Aquilegia* Species

**DOI:** 10.1101/2025.04.09.647901

**Authors:** Yinmei Ma, Junchu Peng, Yuling Weng, Huiqiong Li, Zhi-Qiang Zhang

## Abstract

Mating systems play a crucial role in plant speciation. In particular, hybrid seed inviability is prevalent among species with different mating systems due to parental conflict for resource allocation to offspring. However, whether such a postzygotic barrier can be rapidly established in recently diverged species remains poorly understood. In this study, we integrate population genomic and ecological approaches to address this question using recently diverged species pairs *Aquilegia kansuensis* and *A. ecalcarata*, which diverged approximately 0.256 Ma, in a sympatric population from Qinghai, China. Population structure and demographic history results reveal clear genetic differentiation between the two species, corroborating their recent divergence. The results of outcrossing rate estimation based on microsatellite markers indicated that *A. kansuensis* and *A. ecalcarata* exhibit high selfing and mixed mating systems, respectively. We performed reciprocal crosses between *A. kansuensis* and *A. ecalcarata* and found that *A. ecalcarata* yielded a very small number of small-sized seeds when selfed *A. kansuensis* was used as pollen donors, resulting in strong asymmetric reproductive isolation. An approximate Bayesian computation framework identified that approximately 4.6% of genomic loci are associated with reproductive isolation, and gene ontology analyses of these loci highlighted key pathways involved in seed maturation and post-embryonic development. Therefore, our findings provide evidence that *A. ecalcarata* and *A. kansuensis* represent a novel case of parental conflict and postzygotic isolation driven by divergent mating systems, suggesting mating systems can play a critical role in rapid plant speciation.

## Introduction

According to the biological species concept, speciation occurs through the gradual establishment of reproductive isolation barriers. Understanding the evolution of these barriers is, therefore, critical to understanding speciation processes (Coyne and Orr 2004). Typically, gene flow between diverging lineages is reduced by multiple barriers that act sequentially before zygote formation or post-zygotically (Coyne and Orr 2004; Baack et al 2015). In the traditional view, speciation generally begins with geographic isolation, during which natural selection drives genetic divergence, leading to hybrid incompatibility and ecological differentiation (Coyne and Orr 2004). However, species can also differentiate rapidly in the absence of strong geographic barriers, driven by environmental factors and ecological pressures (McGee et al. 2020, Turbek et al. 2021). Understanding the key reproductive barriers and the underlying mechanisms driving their evolution has been a major goal of rapid speciation. Yet, this remains a challenging task.

The mating system refers to the pattern and frequency of mating among individuals within a population (Barrett and Harder 2017). Studies on animals indicate that selective mating patterns based on preference can quickly form pre-mating isolation without geographic isolation, making it a critical mechanism for rapid speciation (Seehausen et al. 2008, Turbek et al. 2021). Although most flowering plants are hermaphroditic and primarily rely on animal pollination for sexual reproduction (Tong et al. 2023), the mating systems among species and populations exhibit a wide diversity ranging from predominantly selfing to strictly outcrossing (Goodwillie et al. 2010, Whitehead et al. 2018). Plant mating systems could serve as crucial reproductive barriers among closely related plant taxa in several ways (Cutter 2019; Pickup et al. 2019; Gorman et al. 2021; Marie-Orleach et al. 2024).

Firstly, mating systems can directly influence the establishment of prezygotic reproductive barriers. Self-fertilization reduces gene flow from heterospecifics by giving self-pollen a competitive advantage in ovule fertilization when it reaches the stigma before heterospecific pollen, thereby forming a significant prezygotic barrier (Brys et al. 2016). In addition, selfing leads to the rapid evolution of smaller flowers, reduced dichogamy or herkogamy, and lower pollen production in selfing lineages—a phenomenon known as the “selfing syndrome” (Sicard and Lenhard 2011, Slotte et al. 2013, De Vos et al. 2014, Kofler et al. 2024). The syndrome may influence pollinator behavior and promote reproductive isolation between selfing lineages and their outcrossing ancestors (Cutter, 2019).

Furthermore, mating systems can also contribute to postzygotic reproductive isolation by influencing the strength of parental conflict. Theory predicts that outcrossing is thought to increase the intensity of parental conflict while inbreeding is thought to reduce it; outcrossing parents, therefore, are expected to “overpower” selfing parents in crosses between plants with differing mating(Brandvain and Haig 2005, Lafon-Placette and Köhler 2016). Consequently, the dosage imbalance is expected in the hybrid endosperm, causing nonreciprocal seed defects. Accumulating evidence has revealed that when the selfing species serves as the maternal parent, the paternal genome from the outcrossing species regulates resource uptake, thereby permitting normal seed development (Brandvain and Haig 2005, Martin and Willis 2007, Briscoe Runquist et al. 2014, Brys et al. 2014, Iİltaş et al. 2021, Rifkin et al. 2023, Lan et al. 2024). Consequently, hybrid seed failure acts as a poszygotic barrier that restricts gene flow primarily from selfing to outcrossing lineages (Brandvain and Haig 2005, Lafon-Placette and Köhler 2016, Oneal et al. 2016, Städler et al. 2021, Coughlan 2023). Although prezygotic barriers have traditionally been viewed as the primary drivers of speciation in angiosperms (Baack et al. 2015, Christie et al. 2022), recent evidence underscores the role of the mating system in the evolution of reproductive isolation during rapid plant speciation (Coughlan 2023). However, a recent study on North American Arabidopsis lyrata found that the transition to selfing has not led to hybrid seed failure between selfing and outcrossing populations (Gorman et al 2021). It suggests that the genetic basis of hybrid seed inviability is needed to clarify the role of mating systems in the rapid establishment of species barriers in plants.

The genus *Aquilegia* (Ranunculaceae) comprises approximately 70 species distributed predominantly across the temperate regions of the Northern Hemisphere, making it a new model of adaptive radiation in plants (Kramer 2009, Fior et al. 2013). Current evidence suggests that the *Aquilegia* originated in East Asia during the Late Miocene, with subsequent migration and rapid diversification into Eurasia and North America occurring in the mid-Pliocene (Fior et al. 2013). Within the Eurasian clade, populations found in central and western China form a well-supported monophyletic group (Fior et al. 2013, Huang et al. 2018, Liu et al. 2025). Thses lineages were hypothesized to have migrated from the Taihang Mountains in North China toward the southwestern mountainous regions around 2.91 Ma, followed by a burst of rapid diversification approximately 1 Ma—likely driven by climatic fluctuations (Fior et al. 2013, Xue et al. 2021, Liu et al. 2025). Our preliminary whole-genome resequencing analyses have revealed complex reticulate evolution within this group, with evidence suggesting that two to three cryptic species exist within a single morphologically defined species (submitted to Communications Biology). Notably, the lineages of *A. kansuensis* and *A. ecalcarata* occurring in the Gansu and Qinghai Provinces of China display pronounced differences in floral traits, plant stature, leaf dimensions, and overall flower morphology and coloration (Fig. 1). Despite their highly overlapping distributions and frequent sympatry (Weng et al. 2023), suspected hybrid individuals are rarely observed. Furthermore, based on our field observations over the past five years, we found that in *A. kansuensis* with long-spur phenotype, the anther-stigma distance remains minimal from the first day of flowering, whereas in spurless *A. ecalcarata*, this distance progressively decreases with flower development. We, therefore, hypothesize that contrary to expectations based on traditional selfing syndrome characteristics, *A. kansuensis* predominantly employs selfing while *A. ecalcarata* may primarily rely on outcrossing. Additional ecological and genetic investigations are necessary to confirm these divergent mating systems and to elucidate their contributions to reproductive isolation. Given their significant mating system divergence and recent rapid speciation, sympatric populations between *A. kansuensis* and *A. ecalcarata* might offer an ideal model for exploring the role of mating systems in promoting rapid speciation.

**Figure 1.**
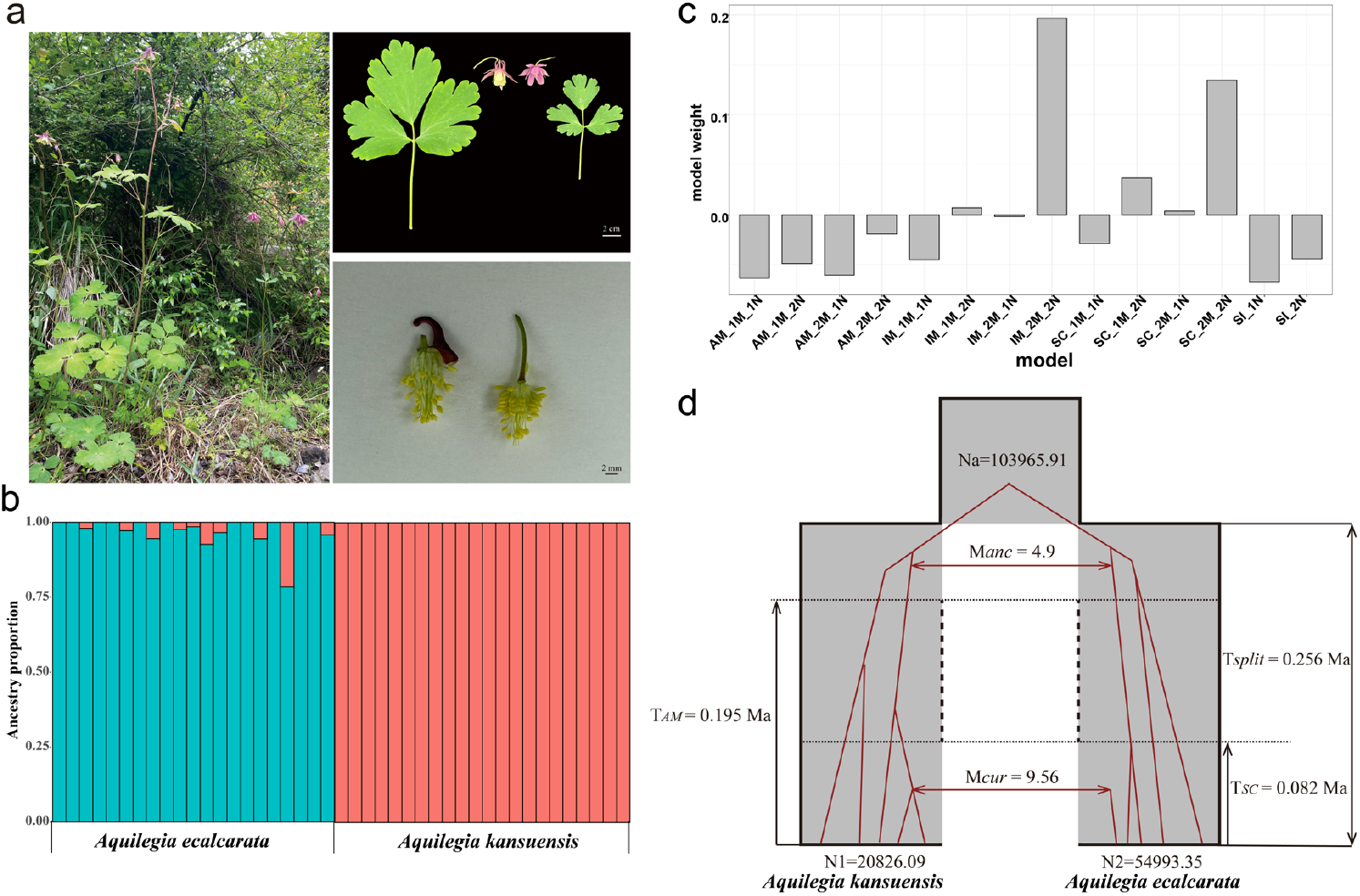
The population structure and demographic history of sympatric population pairs between *Aquilegia ecalcarata* and *Aquilegia kansuensis*. (a) The field photographs of the sympatric population, with *A. kansuensis* individuals on the left and *A. ecalcarata* individuals on the right. (b) Genetic structure analysis at K = 2. (c) Model weights of 14 demographic×genomic models. (d) Hypermodel inferred by RIDGE software based on the weights of 14 models. SI denotes the strict isolation model, SC the secondary contacts model, AM the ancestral migration model, and IM the isolation-migration model. 1M, 2M, 1N, and 2N represent homogeneous or heterogeneous migration rates and effective population sizes, respectively.

## Method and Materials

### Sampling, sequencing, and population structure

Here, we mainly focus on a sympatric population between *A. ecalcarata* and *A. kansuensis* in Qinghai province, China (36.93°N,101.70°E). Besides 20 individuals from our preliminary study (submitted to Communications Biology), we further collected 14 individuals of *A. kansuensis* and 10 individuals of *A. ecalcarata* from this sympatric site. A total of 22 individuals for each morphospecies from this sympatric site (Table S1). Libraries were subsequently sequenced on an Illumina NovaSeq6000 platform at Genedenovo Company (Guangzhou, China). To ensure robust variant calling, the raw reads were stringently filtered to generate high-quality data by: 1) discarding reads containing ≥10% unidentified nucleotides (N); 2) removing reads in which over 50% of the bases exhibited a phred quality score ≤ 20; and 3) excluding reads that aligned to barcode adapter sequences.

BWA mem (Li 2013) from bwa-gtz software (https://github.com/Genetalks/gtz) was used to map the cleaned reads to the *A. kansuensis* genome (Xie et al. 2020). SAMtools and Picard were then used to sort the alignments and remove the duplicate reads (Danecek et al. 2021). Variant calling was performed using HaplotypeCaller and GenotypeGVCFs from GATK v4.2 (McKenna et al. 2010). Finally, loci exhibiting an average coverage depth of less than 3 or greater than 150, or missing rate over 10% were excluded using VCFtools v.0.1.16 (Danecek et al. 2011). The PLINK v1.90b6.21 (Purcell et al. 2007) was employed to generate the unlinked dataset with the flag “--indep-pairwise 50 10 0.2”. Finally, Admixture v1.3.0 (Alexander et al. 2009) was used to estimate genetic structure at K = 2 for the sympatric population, thereby inferring the proportion of introgression between the two morphospecies.

### Inference of demographic history and reproductive barriers

To infer the reproductive isolation loci between *A. ecalcarata* and *A. kansuensis*, we employed the RIDGE software—based on an Approximate Bayesian Computation (ABC) framework, to quantify and identify both the proportion of reproductive barriers and the associated genomic regions between the two species (Burban et al. 2024). First, the RIDGE.sh script is used to generate the prior_bound_suggestion.txt file, which is then utilized to set the prior parameter range for the target SNP dataset. Next, the posterior distributions of several genetic parameters—such as genetic diversity and genetic differentiation—for the target dataset are inferred, and the weights of 14 demographic×genomic models within the RIDGE software are evaluated. When the goodness-of-fit of the posteriors is less than 0.05, the prior bounds are adjusted accordingly. Following this, a hypermodel is generated based on the weights of each model. Furthermore, a random forest algorithm is employed to calculate the Bayes Factor for each locus, along with the associated posterior probabilities linked to reproductive isolation. The reproductive isolation sites are filtered using thresholds of posterior probability greater than 0.5 and Bayes Factor exceeding 50. Finally, the gene ontology (GO) annotation analysis of the putative barrier loci was conducted using KOBAS (Xie et al. 2011).

### Mating system estimation

To assess whether our genomic data supported the inferred divergence in mating systems between *A. kansuensis* and *A. ecalcarata*, we first estimated runs of homozygosity (ROH). Using the whole-genome dataset, we computed ROH in 100-kb windows with PLINK v1.90b6.21 (Purcell et al. 2007). An ROH was defined as an unbroken run of a minimum of 35 homozygous SNPs at 100kb windows. The detected ROH were subsequently categorized by length into small (100–200 kb), medium (200–500 kb), and large (>500 kb) bins.

Moreover, 22 randomly selected individuals per species were further marked during their peak flowering periods. At fruit maturity, fresh young leaves and three fruits were collected from each plant. All samples were preserved with unique identifiers to ensure a one-to-one correspondence between fruit and leaf samples. Leaf samples were preserved by silica gel desiccation to serve as the source of parental material for genetic analysis. Fruits were stored at -20°C to preserve seed viability for subsequent germination experiments. Seeds were subjected to vernalization treatment, and 20 seeds selected from three fruits of each plant were germinated in a constant-temperature incubator at 25°C. Seedlings that successfully germinated served as the progeny source for genetic analysis. DNA was extracted from progeny samples (HiPure DNA Nano Kit D3120, Magen, Shanghai) and parental samples (Plant Genomic DNA Kit DP305, TIANGEN, Beijing) using distinct commercial kits optimized for differential tissue types. Fluorescent PCR amplification was performed using 14 microsatellite loci on DNA from 22 parental plants and 169 offspring seedlings of *A*.*kansuensis*, as well as 19 parental plants and 150 offspring seedlings of *A*.*ecalcarata*. Primers for each locus were labeled with self-developed primers from our research group (See Table S2: FR10, FR51, FR61, FR62, FR63, FR64, FR66, FR68, RF70, FR73, FR75, FR76, FR78, FR79). And sequenced on the ABI 3730 XL sequencing platform (Applied Biosystems). Allele sizes were determined using GENEMAPPER 4.0 (Applied Biosystems), with manual verification of their accuracy. Outcrossing rates were estimated using MLTR version 3.4 (Ritland 2002). The corresponding parameter settings were as follows: resampling was performed at the family level, and standard deviations were calculated based on 1000 independent replicates. Multilocus outcrossing rates (*t*_m_) of two species were obtained.

### Artificial cross-pollination and hybrid-pollination

To investigate whether the two species exhibit the Weak Inbreeder/Strong Outbreeder (WISO) hypothesis (Brandvain and Haig 2005) — which states that in interspecific hybridization between plants with divergent mating systemswhen the outcrosser serves as the maternal parent, symptoms of maternal excess (e.g., smaller seeds) are expected; conversely, in the reciprocal cross, symptoms of paternal excess (e.g., larger seeds) should manifest — artificial pollination experiments were conducted using 21 individuals per species in 2019 and 31 individuals per species in 2020. Before flowering, 1-3 flowers on each plant were emasculated and bagged to prevent autonomous pollination. Once the flowers opened and the stigmas became receptive, mature pollen was collected from at least three plants located ≥ 10 meters away, thoroughly mixed, and used for artificial cross-pollination and interspecific pollination. After pollination, bags were maintained for approximately 45 days until fruit maturation. Mature fruits were then collected, the number of filled seeds in each fruit was counted, and the total seed weight per fruit was measured. Finally, single-weight was calculated by dividing the total seed weight by the number of filled seeds.

We then tested the effects of treatment (intraspecific vs. interspecific), year (2019 and 2020), and their interactions on seeds per fruit of *A. kansuensis* and *A. ecalcarata* by using the zero-inflated generalized linear mixed effects model (GlmmTMB) due to excess zeroes. Employing the glmmTMB package in R (Brooks et al. 2017). In this model, we treated “treatment”, “year” and their interactions as fixed factors, and “plant” as random factors, the seed number of fruit as a response variable with ziGamma distribution (log-link). We tested the single seed weight of the two species across different treatments by using a generalized linear mixed effects model (GLMM). In this model, we treated “treatment” and “species” and their interaction as fixed factors, and “plant” as a random factor, and single seed weight as response variables with Poisson distribution (log-link).

## Results

### Population structure and Demographic history

We further sequenced 24 individuals in this study, generating 186.21 GB of raw data with an average sequencing depth of 14.57X (Table S1). After mapping the reads to the reference genome, we identified 2,509,067 genome-wide SNPs and further extracted a dataset of 326,256 unlinked SNPs for downstream analyses. Population structure analysis (K = 2) revealed that individuals clustered into two distinct groups corresponding to their morphospecies (Fig. 1a and 1b). Notably, ten individuals of *Aquilegia ecalcarata* exhibited shared ancestry with *Aquilegia kansuensis*, whereas no individuals of *A. kansuensis* showed evidence of introgression from *A. ecalcarata*. The proportion of shared ancestry among the admixed *A. ecalcarata* individuals ranged from 1.09% to 20.93% (Fig. 1b).

Among the weights of 14 models describing the history of sympatric populations, the isolation-by-migration (IM) model received the highest weight, followed by the secondary contact (SC) model (Fig. 1c). Moreover, both models exhibit heterogeneity in migration and effective population sizes (2M and 2N, respectively; Fig. 1c). Hypermodel further estimate that divergence between sympatric pairs occurred approximately 0.256 Ma (Tsplit). Initially, ancestral migration was moderate (Manc = 4.9) until it ceased around 0.195 Ma, after which current migration resumed at about 0.082 Ma with a higher rate (Mcur = 9.56; Fig. 1d). These findings suggest that the sympatric populations in Qinghai are the result of recent rapid differentiation, characterized by a distinct genetic structure and limited historical gene flow.

## Determination of mating system types

Inbreeding increases homozygosity, resulting in extended stretches of consecutive homozygous genomic regions, known as runs of homozygosity (ROH) (Gibson et al. 2006). In *A. ecalcarata*, ROH segments are predominantly small (100–200 kb) or medium-sized (200–500 kb), with fewer than 210 and 50 segments, respectively. Large ROH segments (>500 kb) are nearly absent (Fig. 2). In contrast, *A. kansuensis* exhibits significantly higher numbers of ROH segments, with more than 450 small, over 200 medium, and between 10 and 50 large segments (*P* < 0.001; Fig. 2). These findings suggest that *A. kansuensis* is more prone to selfing, resulting in the accumulation of a greater number of long ROH segments over a relatively short divergence time.

**Figure 2.**
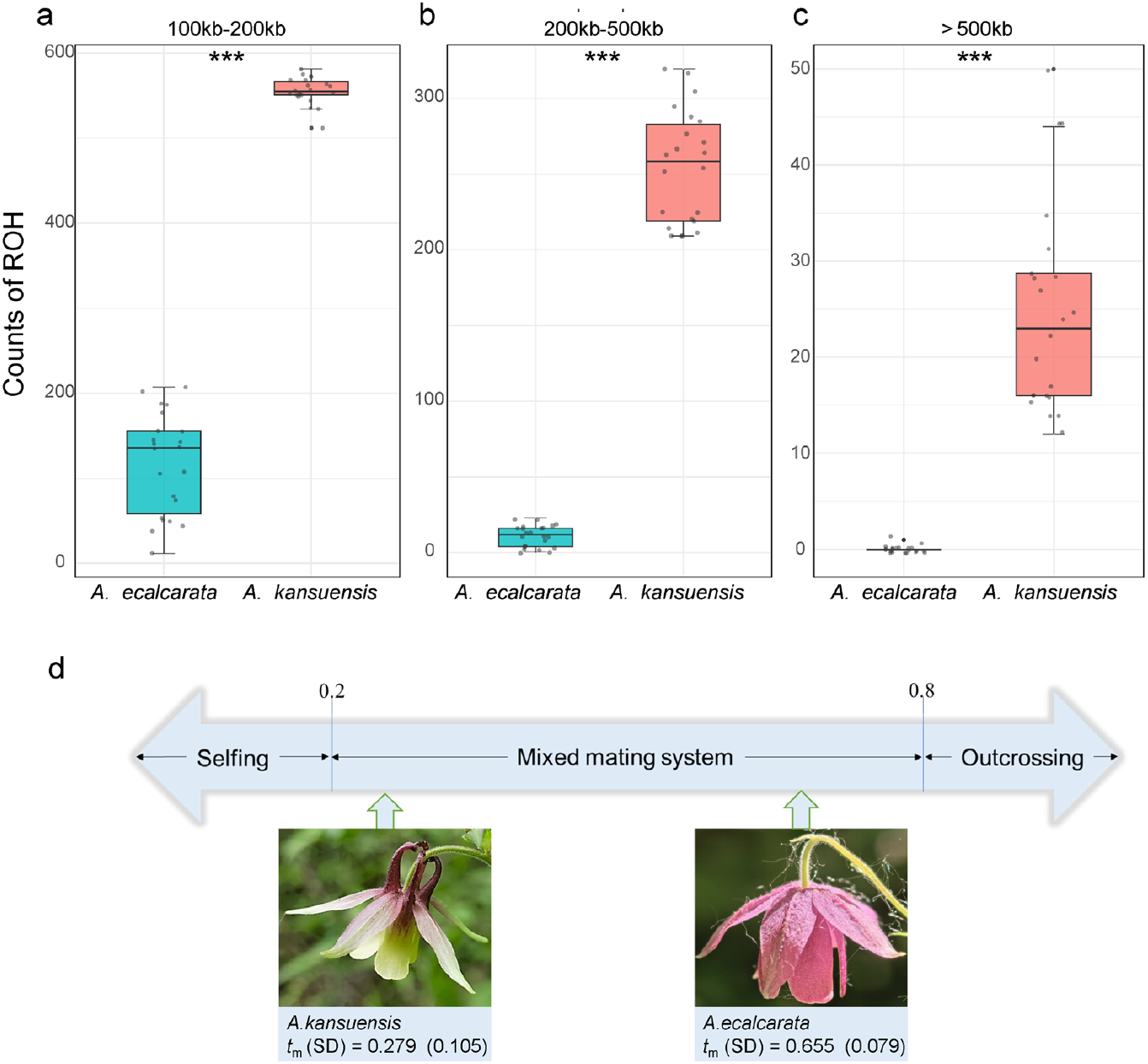
Comparison of mating systems between *Aquilegia ecalcarata* and *Aquilegia kansuensis*. (a-c) Differences in the number of runs of homozygosity (ROH) between *A. ecalcarata* and *A. kansuensis* in sympatric populations with differing mating systems. ROH were categorized into three length classes: small (100–200 kb; panel a), medium (200–500 kb; panel b), and large (> 500 kb; panel c). Across all length categories, *A. kansuensis* exhibited significantly higher ROH counts compared to *A. ecalcarata* (*** represents *P* < 0.001). (d) Multilocus outcrossing rates (*t*_*m*_) for sympatric population pairs, with standard deviations (SD) shown in parentheses.

Based on the estimation from MLTR version 3.4 (Ritland 2002), the multilocus outcrossing rate (SD) of *A. kansuensis* was 0.279 (0.105), while that of *A. ecalcarata* was 0.655 (0.079). According to the criteria of Schemske and Lande (1985) (selfing when tm ≤ 0.2, mixed mating when tm is between 0.2-0.8, and outcrossing when tm > 0.8), both species exhibit a mixed mating system. Therefore, *A. kansuensis* leans toward selfing, whereas *A. ecalcarata* is more inclined to outcrossing.

### Quantity and quality of seeds from cross-pollination and hybrid-pollination

The seed number per fruit in *A. ecalcarata* showed highly significant differences among pollination treatments, but no significant differences across years and in the interaction between years and pollination treatments (Table 1). In both years, the seed number per fruit from intraspecific outcrossing was significantly higher than that from interspecific hybridization in *A. ecalcarata* (Fig. 3a; *P*_2019_ < 0.0001; *P*_2020_ < 0.0001). The seed number per fruit in *A. kansuensis* exhibited highly significant differences across years, but no significant differences among pollination treatments or in their interaction with years (Table 1). In both years, no significant difference was observed between the seed number per fruit from intraspecific outcrossing and interspecific hybridization in *A. kansuensis* (Fig. 3b; *P*_2019_ = 0.7870; *P*_2020_ = 0.9881). Single seed weight showed highly significant differences among species, treatments, and their interactions (Table 1). In *A. ecalcarata*, single seed weight from intraspecific outcrossing was significantly higher than that from interspecific hybridization (Fig. 4a and 4b; *P* = 0.0024), whereas in *A. kansuensis*, interspecific hybridization resulted in significantly higher single seed weight compared to intraspecific outcrossing (Fig. 4a and 4c; *P* = 0.0071).

**Table 1.** Results of Zero-inflated Generalized Linear Mixed Effects model (GlmmTMB) analyzing differences in the seed number of fruit across pollination treatments and years, and Generalized Linear Mixed Models (GLMM) analyzing differences in single seed weight between species and pollination treatments.

| Sources of variation | Chisq | Df | *P* |
| --- | --- | --- | --- |
| ***A. ecalcarata*: Seed number of fruit** |  |  |  |
| Treatment | 186.4601 | 1 | <0.001 |
| Year | 0.1499 | 1 | 0.6986 |
| Treatment : Year | 0.9292 | 1 | 0.3351 |
| ***A. kansuensis*: Seed number of fruit** |  |  |  |
| Treatment | 0.6658 | 1 | 0.4145 |
| Year | 25.8085 | 1 | <0.001 |
| Treatment : Year | 0.3118 | 1 | 0.5766 |
| **Single seed weight** |  |  |  |
| Treatment | 13.197 | 1 | <0.001 |
| Speices | 301.379 | 1 | <0.001 |
| Speices : Treatment | 21.964 | 1 | <0.001 |

**Figure 3.**
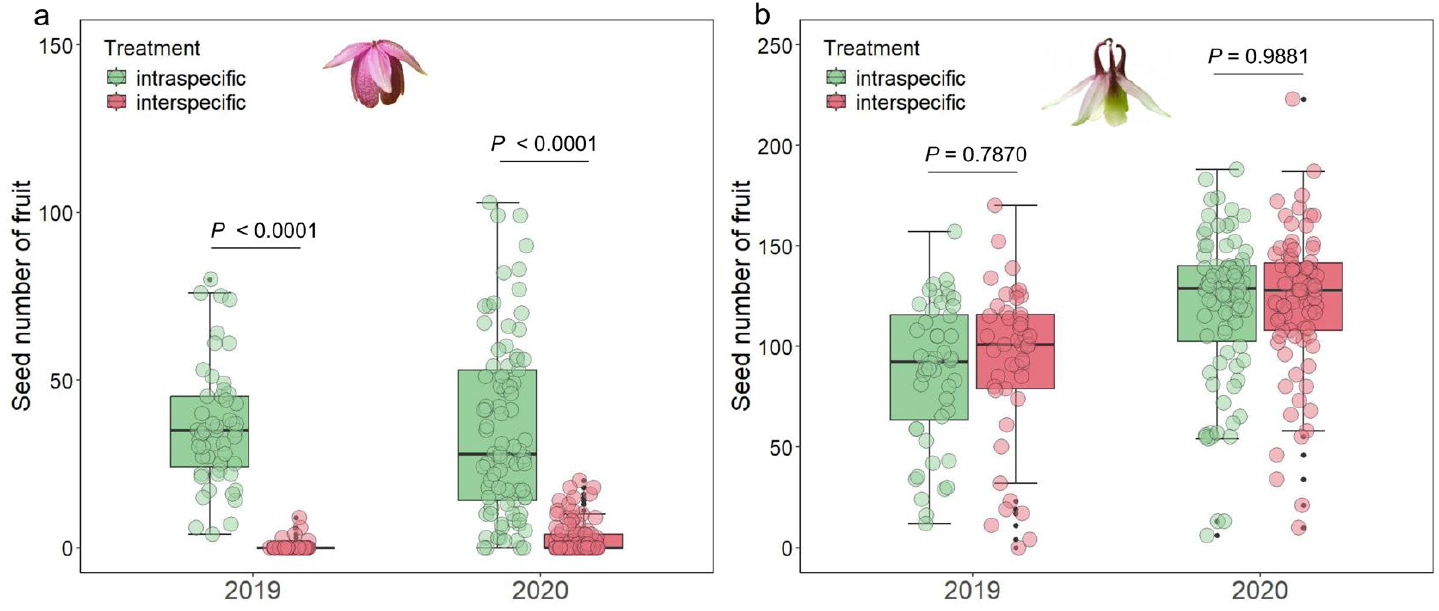
Seed number per fruit from artificial intraspecific outcrossing and interspecific hybridization pollination in two species across different years. (a) the seed number of fruit following two pollination treatments conducted over two years with *A*.*ecalcarata* as the maternal parent, (b) the seed number of fruit following two pollination treatments conducted over two years with *A. kansuensis* as the maternal parent.

**Figure 4.**
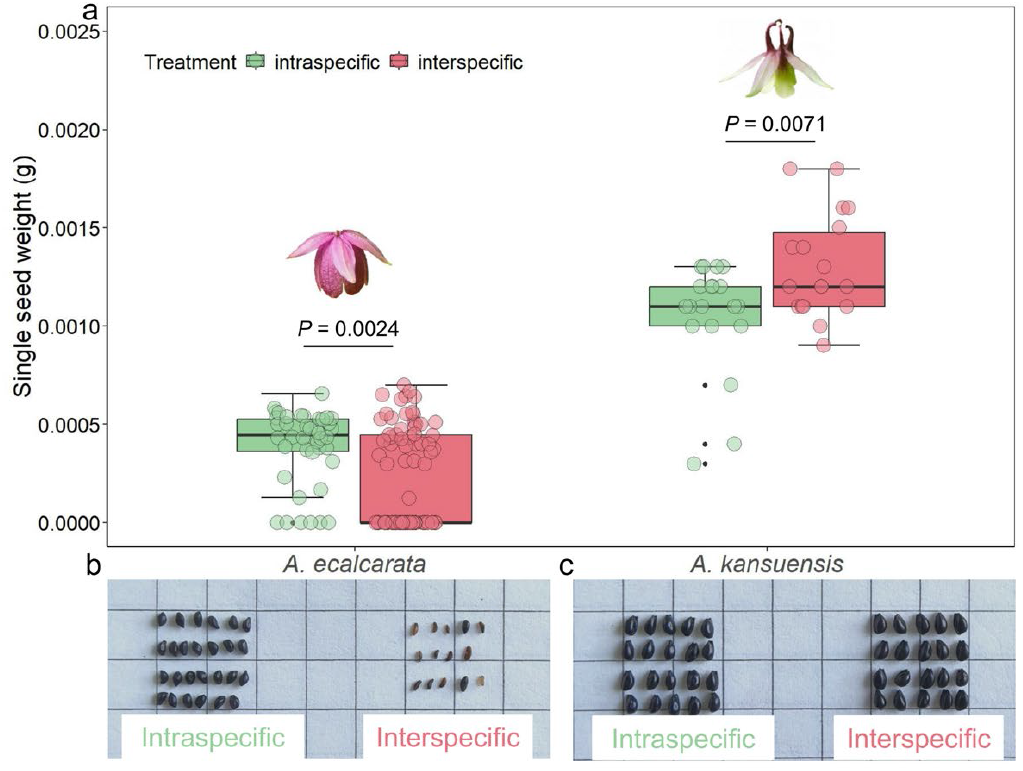
Single seed weight from artificial intraspecific outcrossing and interspecific hybridization pollination in two species. a: the single seed weight following two pollination treatments with *A*.*ecalcarata* as the maternal parent, b: the single seed weight following two pollination treatments with *A*.*kansuensis* as the maternal parent.

### Reproductive isolation sites and GO annotation

Based on the hypermodel from RIDGE software, we further screened the reproductive isolation sites between *A. kansuensis* and *A. ecalcarata*. Our results revealed that 4.6% of the 50-kb windows (255 out of 5518; Table S3) exhibited posterior probabilities greater than 0.5 and Bayes factors exceeding 50, and these regions were classified as barrier loci (Table S3). The genes associated with these barrier loci were subsequently annotated (Table S4). Notably, genes involved in post-embryonic development, pollen maturation, and the positive regulation of seed germination were identified (Table S4), which may further support the role of mating system differences in driving reproductive isolation between *A. ecalcarata* and *A. kansuensis*.

## Discussion

This study focused on mating systems and their role in postzygotic reproductive isolation between sympatric *A. kansuensis* and *A. ecalcarata*. By integrating population genetics with field-based reproductive ecology, we evaluated the mating systems and post-fertilization reproductive isolation between these two recently divergent species. Our results indicate that the mating system might play a critical role in the rapid evolution of postzygotic reproductive isolation.

Although the divergence time between the two species is relatively short, around 0.256 Ma (Fig. 1d), they exhibited significant differences in floral morphology, plant height, leaf size, and flower coloration (Fig. 1a). with rare putative hybrids in the sympatric populations (our observation). Moreover, the high degree of range overlap and frequent sympatric distribution (Weng et al. 2023) indicate a lack of pronounced geographic isolation. However, the genetic structure and demographic history revealed clear genetic differentiation and low historical gene flow between the two species (Figs. 1b and 1d), suggesting that a high degree of reproductive isolation has been established, effectively limiting gene exchange among sympatric populations. Predictions from the RIDGE model also support that an isolation-with-migration (IM) model without geographic isolation best explains the formation of these sympatric populations (Fig. 1c). Additionally, the hypermodel that incorporates a secondary contact scenario reveals that historical gene flow did not significantly decrease after a possible phase of geographic isolation; rather, it appears to have increased (Fig. 1d). These findings support that geographic and climatic factors are not the primary drivers of divergence and the accumulation of reproductive isolation between *A. kansuensis* and *A. ecalcarata* in the Qinghai region.

In contexts where clear geographic and ecological differentiation is lacking, rapid speciation is often closely associated with sexual selection or shifts in mating systems. For instance, the rapid speciation observed in seedeaters from Iberá is tightly linked to mate choice (Turbek et al. 2021). In plant speciation processes devoid of geographic isolation, both pollinator interactions and mating system dynamics can be pivotal. While rapid adaptive radiation in North American *Aquilegia* species has been driven by pollinators shift (Kramer 2009), no significant differentiation in pollinators has been observed in *Aquilegia* species in western China (Tang et al. 2007, Weng 2021, Toji et al., 2022). These results raise the question: how do they remain sufficiently reproductively isolated in sympatric populations?

Our preliminary observations revealed that pollinators exhibiting greater fidelity to the *A. ecalcarata* during interspecific foraging transitions (Weng 2021). However, this behavioral preference alone was unable to completely impede gene flow. The transition between self-fertilization and outcrossing can lead to a series of genetic consequences at the genomic level, such as selfing species retaining longer homozygous segments and exhibiting lower heterozygosity (Curik et al. 2014, Ceballos et al. 2018). Based on SSR molecular markers and whole-genome ROH estimations, significant differences in the homozygous segments and heterozygosity were observed between *A. kansuensis* and *A. ecalcarata*. Contrary to our expectations, *A. kansuensis*, which possesses spur and floral nectar, is a highly self-fertilizing species (*t*_m_ (SD) = 0.279 (0.105)), and has accumulated numerous long ROH segments; in contrast, *A. ecalcarata*, lacking spur and floral nectar, exhibits a predominantly outcrossing mixed mating system (*t*_m_ (SD) = 0.655(0.079)), accumulating only a few short ROH segments (Fig. 2). Therefore, our results support that a shift in the mating system occurred within a relatively short period in *A. ecalcarata* and *A. kansuensis*. We also observed *A. kansuensis* lacks spatial segregation between stigma and anther at the early stage of anthesis (Fig1 a), suggest a possibility of prior selfing or competing selfing in *A. kansuensis*. Thus, we predict that autonomous selfing may act as an important prezygotic reproductive barrier (Brys et al 2016).

All *Aquilegia* species are diploid, self-compatible, and most show high interspecific cross-compatibility with minimal reproductive barriers (Kramer 2009). Even artificial crosses between species endemic to North America and China have yielded viable hybrids (Ballerini et al. 2020). However, our artificial pollination experiments revealed a strong unidirectional hybrid incompatibility between *A. ecalcarata* and *A. kansuensis* (Fig. 3 and Fig. 4). Specifically, when *A. kansuensis* acted as the pollen donor, the vast majority of resulting seeds failed to develop normally. The result consist with the WISO hypothesis advanced by Brandvain and Haig (2005), outcrossing plant species exhibit a more pronounced parental influence on endosperm development—quantified as the Endosperm Balance Number (EBN)—than selfing species, due to stronger selective pressures favoring paternal resource allocation and resisting maternal constraints. It suggest divergent mating systems influence the the detrimental effects of parent conflict on endosperm development and ultimately resulting in seed lethality (Brandvain and Haig 2005, Rebernig et al. 2015, Coughlan 2023). Few plants have also been documented that rapid divergence between a selfing and an outcrossing species, which led to EBN divergence and hybrid seed lethality, like *Arabidopsis lyrate, Capsella grandiflora* and *Capsella rubella* (Guo et al. 2009, Rebernig et al. 2015, İltaş et al. 2021). Furthermore, our results identified several key pathways, including post-embryonic development (GO:0009791), seed development (GO:0048316), and seed maturation (GO:0010431), among others, that may contribute to postzygotic isolation and the regulation of seed maturation (Table S4).

For example, Aqoxy7G02817 is homologous to UmamiT14 (TAIR: At2g39510; Table S4) in *Arabidopsis thaliana*; loss-of-function mutants of this gene accumulate high levels of free amino acids in fruits and produce smaller seeds. Therefore, our findings support that *A. ecalcarata* and *A. kansuensis* represent a new case of parental conflict and postzygotic isolation induced by a rapid transition in mating systems. Thus, our results suggest mating system might be one of the major determinants of strong asymmetric postzygotic reproductive isolation between recently diverged *A. ecalcarata* and *A. kansuensis*.

In addition to the postzygotic isolation mechanisms explored in this study, selfing contributes to reproductive isolation through multiple pathways. First, the selfing syndrome drives floral morphological changes (e.g., reduced anther-stigma distance), which significantly diminish opportunities for cross-pollination (Barrett et al. 1996, Brys et al. 2016). Second, altered gamete interaction mechanisms—such as S-RNase-mediated recognition and rejection of heterospecific pollen—enhance the ability of selfing lineages to filter out foreign pollen (Kubo et al. 2010, Rifkin et al. 2023). Third, frequent genomic structural variations (e.g., inversions/translocations) in selfing lineages directly block gamete fusion during hybridization via Dobzhansky-Muller incompatibilities (Lynch and Force 2000, Taylor et al. 2001). These mechanisms collectively reinforce reproductive barriers at the prezygotic stage.

Furthermore, in selfing populations, genes associated with male reproductive functions may gradually lose functionality or even be deleted due to relaxed sexual selection pressure (Hamlin et al. 2017). This functional divergence not only disrupts reproductive traits but may also affect other linked genes through genetic linkage effects, thereby exacerbating genetic divergence between populations at the molecular level (Li et al. 2023). Therefore, the co-evolution of the selfing syndrome and genomic structural variation may constitute a “pre-adaptation–isolation– differentiation” cascade. This hypothesis requires further validation through comparative genomics and experimental evolution approaches. Understanding these mechanisms holds not only theoretical significance but also practical implications for the hybridization management of endangered species and the improvement of crop domestication.

## Acknowledgments

This work was financially supported by funding from the National Natural Science Foundation of China (32271693) and the Cultivating Plan Program for the Leader in Science and Technology of Yunnan Province (202405AC350111).

## Author contributions

ZQZ conceived and developed the research. YM, YW, and HL performed the field investigations and experiments and analyzed the ecological data. JCP analyzed the population genomic data. All authors contributed to writing the first draft and critically reviewed the final manuscript.

